# PPMO-mediated exon skipping induces uniform sarcolemmal dystrophin rescue with dose-dependent restoration of circulating microRNA biomarkers and muscle biophysical properties

**DOI:** 10.1101/2022.01.25.477672

**Authors:** Katarzyna Chwalenia, Jacopo Oieni, Joanna Zemła, Małgorzata Lekka, Nina Ahlskog, Anna M.L. Coenen-Stass, Graham McClorey, Matthew J.A. Wood, Yulia Lomonosova, Thomas C. Roberts

**Affiliations:** Department of Paediatrics, University of Oxford, South Parks Road, Oxford, OX1 3QX, UK; Department of Biophysical Microstructures, Institute of Nuclear Physics, Polish Academy of Sciences, PL-31342 Kraków, Poland; Department of Physiology, Anatomy and Genetics, University of Oxford, South Parks Road, Oxford, OX1 3QX, UK; MDUK Oxford Neuromuscular Centre, South Parks Road, Oxford, OX1 3QX, UK

**Keywords:** Duchenne muscular dystrophy, biomarkers, miRNA, myomiR, extracellular, *mdx*, exon skipping, PPMO

## Abstract

Duchenne muscular dystrophy (DMD) is a paediatric muscle-wasting disorder caused by genetic loss of the gene encoding the dystrophin protein. Therapies that restore dystrophin expression are presumed to correct the disease, with antisense-mediated exon skipping being the leading approach. In this study, we aimed to determine whether exon skipping using a peptide-phosphorodiamidate morpholino oligonucleotide (PPMO) conjugate results in dose-dependent restoration of uniform dystrophin localization, together with correction of putative DMD serum and muscle biomarkers. To this end, dystrophin-deficient *mdx* mice were treated with a PPMO (Pip9b2-PMO) designed to induce *Dmd* exon 23 skipping and dystrophin rescue at single, ascending intravenous doses (3, 6, or 12 mg/kg) and sacrificed two weeks later. Dose-dependent exon skipping and dystrophin protein restoration were observed. Importantly, dystrophin expression was uniformly distributed at the sarcolemma of corrected myofibers at all doses. The abundance of serum microRNA biomarkers (i.e. miR-1a-3p, miR-133a-3p, miR-206-3p, miR-483-3p) and creatinine kinase were restored towards wild-type levels after treatment in a dose-dependent manner. All biomarkers were strongly anti-correlated with both exon skipping level and dystrophin expression. Dystrophin rescue was also strongly positively correlated with muscle stiffness (i.e. Young’s modulus) as determined by atomic force microscopy (AFM) nanoindentation assay. These data demonstrate that PPMO-mediated exon skipping generates myofibers with uniform dystrophin expression, and that both serum miRNA biomarkers and muscle AFM have potential utility as pharmacodynamic biomarkers of dystrophin restoration therapy in the context of DMD.

## Introduction

Duchenne muscular dystrophy (DMD) is an X-linked, monogenic muscle-wasting disorder, and the common inherited myopathy affecting children. DMD is caused by loss-of-function mutations in the gene encoding dystrophin (*DMD*), which performs structural and signalling functions at the sarcolemma ^1–3^. The leading approach for treating DMD is exon skipping, whereby splicing of the *DMD* pre-mRNA is modulated using antisense oligonucleotides, such that the translation reading frame is corrected, and an internally-truncated, partially-functional quasi dystrophin protein produced ^4^. At the time of writing (December 2021) there are four FDA-approved exon skipping drugs for DMD: eteplirsen, viltolarsen, golodirsen, and casimersen (all unconjugated phosphorodiamidate morpholino oligonucleotides (PMO)s targeted for skipping of *DMD* exons 51, 53, 53, and 45, respectively) ^5^. However, the levels of dystrophin achieved in treated DMD patient muscle are very low (∼1% of healthy levels) ^6,7^. Studies of dystrophin restoration therapies have primarily focused on the amount and quality of dystrophin protein expressed ^8^. In contrast, the importance of correct dystrophin localization has been less well-studied. Notably, recent work from our group ^9^, and others ^10^, has suggested that uniform dystrophin expression is an important factor in the success of dystrophin restoration therapies and the severity of dystrophinopathies, respectively. Furthermore, challenges in determining dystrophin expression (e.g. the requirement for invasive biopsy, whether a single biopsy is reflective of the overall musculature, and technical difficulties associated with accurate measurement of dystrophin expression) have motivated the search for alternative DMD biomarkers ^30^. For example, extracellular microRNAs (ex-miRNAs) constitute one such class of minimally-invasive biomarkers that may have utility in the context of DMD ^11^. The leading candidate miRNA biomarkers of DMD are the myomiRs (muscle-enriched miRNAs that regulate myoblast proliferation and differentiation ^12–15^), consisting of miR-1a-3p, miR-133a-3p, and miR-206-3p. These myomiRs are highly upregulated in DMD patient serum ^16,17^ and in various dystrophic mouse models including *mdx* ^16–20^, *mdx52* ^21^, dKO (dystrophin/utrophin double knock-out) ^22^, and *mdx^4cv^* ^23^. In the *mdx* mouse, it has been reported that extracellular myomiRs (ex-myomiRs) are less variable than serum creatine kinase (CK) ^18^, the gold standard clinical chemistry biomarker for neuromuscular disease. Multiple other putative DMD biomarkers have been identified ^24,25^. For example, we recently identified miR-483-3p as being enriched in dystrophic mouse serum ^9,17^. It is currently not clear whether any of these additional miRNA biomarkers offers an advantage over serum myomiRs.

ex-myomiRs offer excellent discrimination between DMD patients and healthy controls ^16,17^, suggesting they could be used to diagnose patients. However, the myomiRs have also been found to be elevated in the serum of multiple other neuromuscular conditions (including myotonic dystrophy, spinal muscular atrophy, and amyotrophic lateral sclerosis) ^23,26–29^ meaning that they are not specific biomarkers for DMD. Importantly, diagnosis of DMD is not currently a major clinical challenge. Instead, these ex-miRNAs may have utility as minimally-invasive biomarkers for monitoring disease progression, or response to therapy, a currently unmet clinical need ^11,30^.

Work from our group in the *mdx* mouse has shown that ex-myomiRs are primarily non-vesicular, and are instead protected from nucleolytic degradation through interactions with serum proteins ^20,31^. We have proposed that extracellular myomiRs are markers of muscle turnover (i.e. myonecrosis and compensatory regeneration) which are elevated during periods of regenerative activity in *mdx* muscle ^20,32^, and are progressively secreted during myogenic differentiation in culture ^32^. Furthermore, there is a degree of selectivity in myomiRs release ^17,32^, suggesting that passive leakage from damaged muscle is insufficient to explain their serum abundance levels.

A single intravenous treatment of the *mdx* mouse with peptide-PMO (PPMO, designed to skip *Dmd* exon 23 and thereby rescue dystrophin expression) leads to restoration of ex-myomiRs towards wild-type levels ^9,17,19,20^. Similarly, ex-myomiR levels in dystrophic mouse serum are restored to near-wild-type levels after treatment with the U1 ^16^ and U7 ^22^ small nuclear RNA expressed exon skipping systems. Importantly, we observed that treatment with two PPMO conjugates with different potencies resulted in different degrees of ex-myomiR restoration ^20^, suggesting that ex-myomiR levels might be suitable for non-invasively assessing dystrophin levels in muscle. We further investigated this phenomenon using a genetic model (the *mdx-Xist*^Δhs^ mouse) in which dystrophin is expressed at variable levels as a consequence of skewed X-chromosome inactivation ^33^. We generated animals that expressed ∼10%, ∼20% and ∼35% of healthy dystrophin protein levels, but surprisingly, ex-myomiR levels were not correlated with dystrophin levels at all, and were instead elevated at similar levels to dystrophin-null *mdx* controls, regardless of the level of dystrophin expression ^9^. Importantly, dystrophin is expressed in a within-fiber patchy manner in the *mdx-Xist*^Δhs^ model ^9,33^, suggesting that a uniform pattern of sarcolemmal dystrophin is important for therapeutic correction. Here, we aimed to determine if PPMO-mediated exon skipping resulted in dose-dependent uniform dystrophin expression, ex-myomiR restoration, and correction of the biophysical properties of dystrophic muscle.

## Results

### Dose-dependent dystrophin restoration in *mdx* muscle following PPMO treatment

Male *mdx* mice were treated with a single intravenous injection (via the tail vein) of PPMO conjugate (Pip9b2-PMO) (**Figure 1A**) at 8 weeks of age. Mice were treated with one of three doses: 3 mg/kg (*n*=7), 6 mg/kg (*n*=9), or 12 mg/kg (*n*=9). These treatments were intended to induce three different levels of dystrophin re-expression corresponding to each dose. Animals were sacrificed two weeks post treatment (i.e. 10 weeks old) at which point blood serum and tibialis anterior (TA) muscles were harvested. Untreated (age- and sex-matched) *mdx* (*n*=9) and wild-type (WT, *n*=5) animals were harvested in parallel as controls (**Figure 1B**). Dose-dependent restoration of dystrophin protein expression was confirmed by western blot with median expression values of 3.7%, 16.7%, and 44.8% of WT levels for the 3, 6, and 12 mg/kg PPMO-treatment groups, respectively (**Figure 1C,D**). Dystrophin protein was undetectable in untreated *mdx* TA muscle lysates (**Figure 1C**). The levels of exon-skipped (ΔExon23) *Dmd* transcripts in PPMO-treated muscle was similarly dose responsive (**Figure 1E**), as determined by RT-qPCR. Median skipped transcript levels were 1.2%, 11.8%, and 38% of total transcripts for the three PPMO-treatment dose groups, respectively. Dystrophin protein quantification and exon skipped transcript levels were strongly correlated (Spearman’s *r*=0.9262, *P*<0.0001, **Figure 1F**).

**Figure 1.**
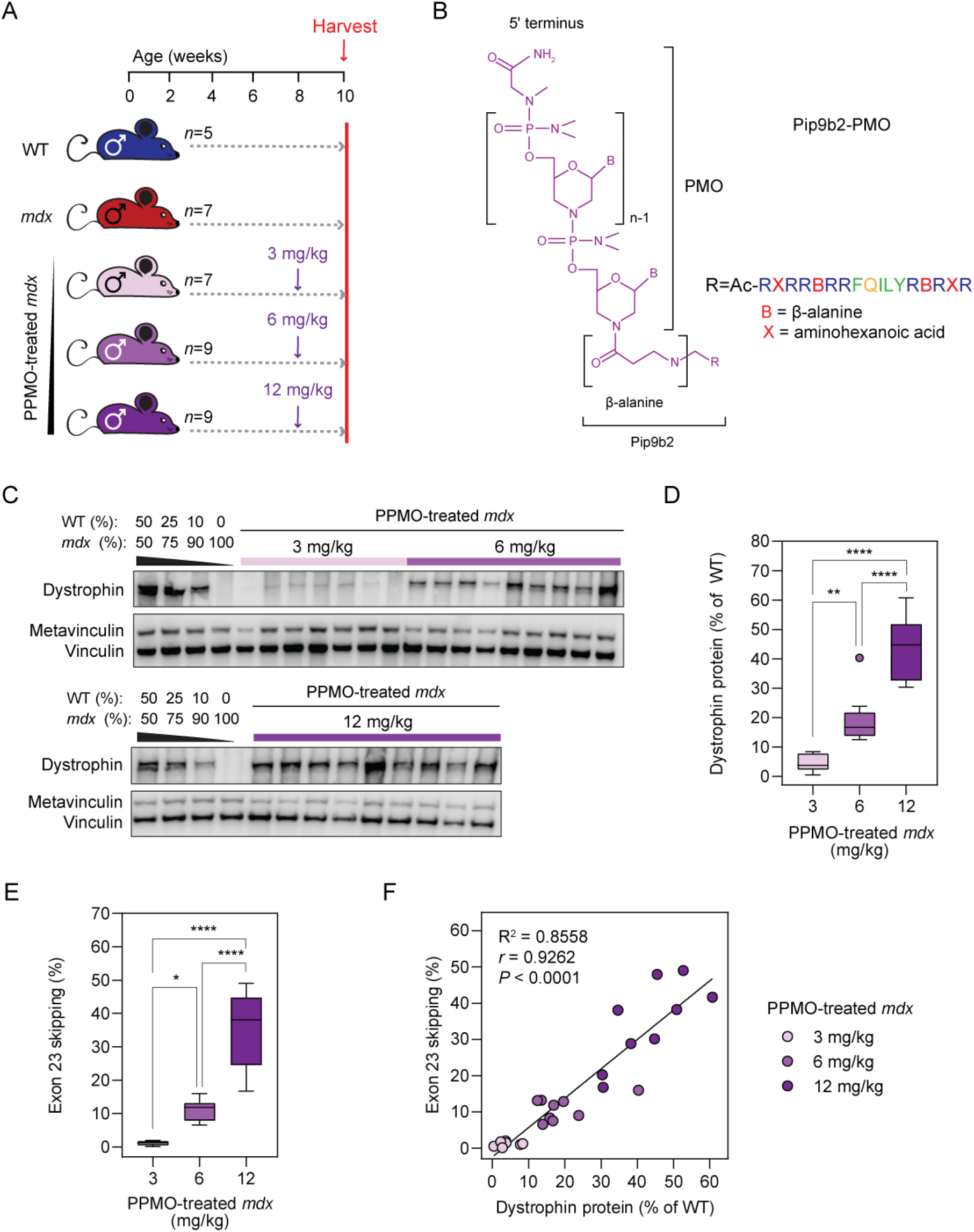
Dose-dependent dystrophin restoration in *mdx* mice treated with PPMO conjugates. (**A**) Schematic of experimental design. Eight-week-old male *mdx* mice were treated with a single intravenous injection of PPMO conjugates (at a dose of 3, 6, or 12 mg/kg) and harvested for analysis two weeks later. (**B**) Structure of the PPMO conjugates, which consist of a Pip9b2 peptide covalently conjugated to a PMO antisense oligonucleotide designed to skip *Dmd* exon 23. (**C**) Western blot of dystrophin protein expression for all PPMO-treated *mdx* TA muscle lysates. Vinculin was used as a loading control. (**D**) Tukey box plot of dystrophin protein expression quantification. (**E**) Tukey box plot of *Dmd* exon 23 skipping as determined by RT-qPCR. Statistical significance was tested by one-way ANOVA and Bonferroni *post hoc* test. \**P*<0.05, \*\**P*<0.01, \*\*\*\**P*<0.0001. (**F**) Correlation plot of dystrophin protein expression and exon 23 skipping levels for PPMO-treated *mdx* samples. Values were analysed using linear regression and Spearman correlation analyses.

### Validation of myomiR RT-qPCR assay specificity

The leading serum miRNA biomarkers in the context of DMD are the myomiRs ^11^, consisting primarily of two miRNA families: (i) the miR-1/206 family (i.e. miR-1a-3p and miR-206-3p), and (ii) the miR-133 family (i.e. miR-133a-3p and miR-133b-3p). miR-1a-3p and miR-206-3p share the same seed sequence and differ only at 4 nucleotides. Conversely, miR-133a-3p and miR-133b-3p are identical except for the 3ʹ terminal nucleotide (**Figure 2A**). Despite their sequence similarity, miR-133a-3p and miR-133b-3p are derived from distinct genomic loci, and are expressed at different levels in murine muscle, with miR-133a-3p being ∼5-fold more abundant than miR-133b-3p (**Figure S1A**) ^17^. The first report of serum myomiR levels in dystrophic serum commented that the RT-qPCR methodology used (miScript, developed by Qiagen) was unable to distinguish between miR-133a-3p and miR-133b-3p ^16^. Since that initial report, several groups have reported serum miRNA quantification data for these two miRNAs in the context of DMD ^23,34–36^, myotonic dystrophy ^29,37–39^, and in human foetal myogenesis ^40^. These studies utilized the small RNA TaqMan RT-qPCR method (from Applied Biosystems, now Thermo Fisher Scientific) ^41^, which involves an RNA-specific stem-loop primer in the reverse transcription (RT) step. The small RNA TaqMan approach is by far the most widely adopted miRNA quantification technology at use in this field (**Figure S2B**). We were therefore motivated to determine if these assays are sufficiently specific to distinguish between miR-1a-3p and miR-206-3p, and between miR-133a-3p and miR-133b-3p. Synthetic, single-stranded RNA oligonucleotide mimics of these four miRNAs were analysed in separate reactions using either the specific assay (i.e. both the stem loop primer, and the TaqMan assay) or with the assay corresponding to the related family-member miRNA. For the miR-1/206 family, the assays were found to be highly specific, with greater than 40,000-fold difference (ΔCq>15.4) between on-target and off-target amplification (**Figure 2B**). Cq values for the off-target amplification were ∼31-35, suggestive of low or very low levels of amplification. Conversely, the miR-133 family assays were not capable of discriminating between miR-133a and miR-133b, as equivalent amplification was observed when the miR-133a and miR-133b oligonucleotides were analysed with either assay (ΔCq=0.3) (**Figure 2C**). This lack of specificity could be attributed to the RT step, as negligible amplification was observed when the miRNA mimics were first reverse transcribed using the correct stem loop primer, followed by qPCR with the off-target TaqMan assay (**Figure S2**). The reason for the cross-reactivity of these assays becomes apparent when the interactions between the stem-loop primers and the on-target/off-target miRNAs are inspected (**Figure 2D,E**). Stem-loop primers recognize their cognate miRNAs through binding of the six 3ʹ terminal nucleotides of the RT primer with six complementary 3ʹ terminal nucleotides of the miRNA, which permits priming of the RT enzyme (**Figure 2D**). For off-target primer interactions for the miR-1/206 family miRNAs, there are either two or three mismatches between the RT primer and the miRNA, including at the most 3ʹ terminal nucleotide, which is unlikely to support priming (**Figure 2D**). Conversely, for the miR-133 family, there is only one mismatch or one wobble base pair between primer and miRNA for the off-target RT primer combinations, respectively. In particular, the most 3ʹ terminal nucleotide is complementary in both cases, which is expected to support priming, and therefore explains the high levels of off-target miR-133 amplification (**Figure 2E**).

**Figure 2.**
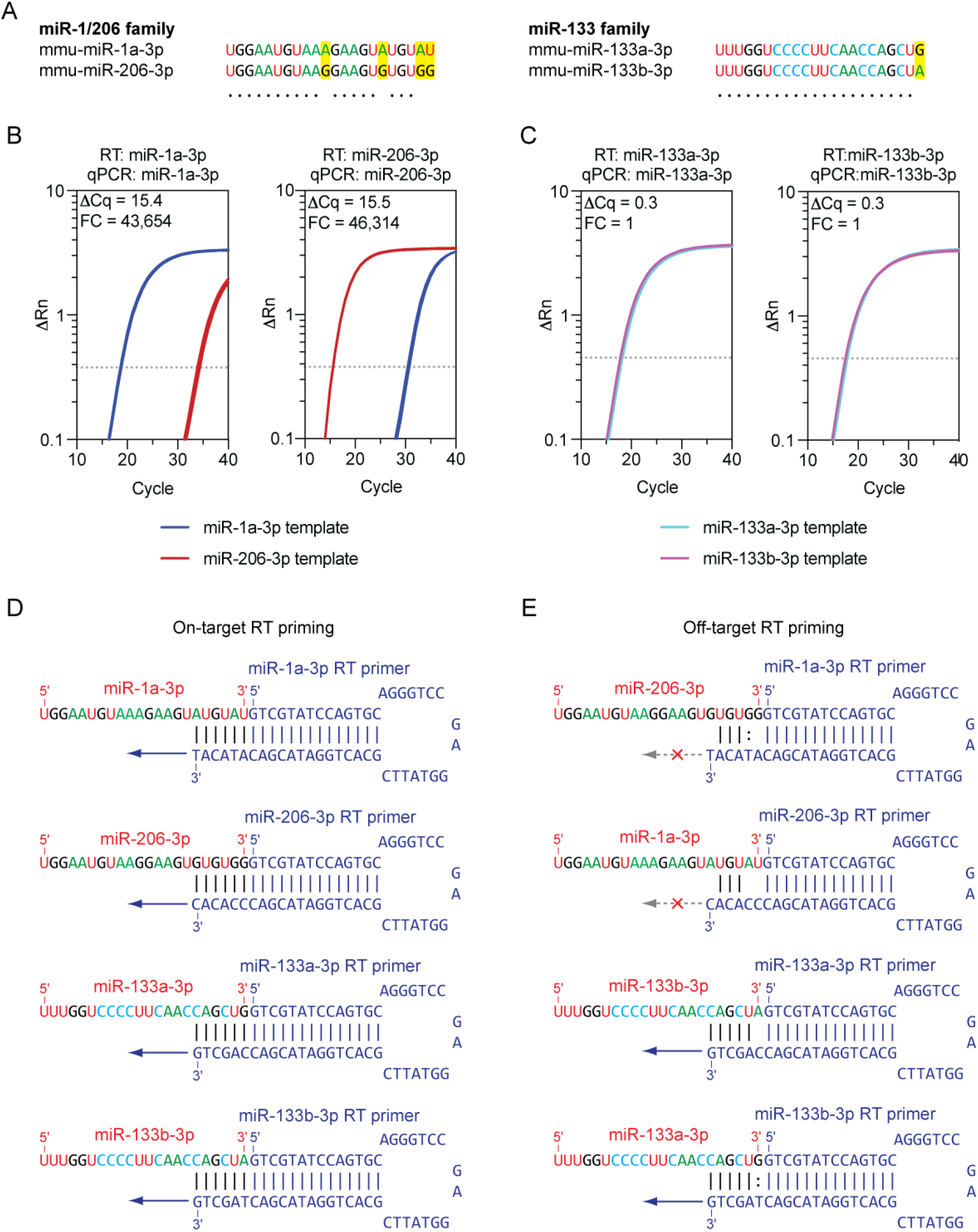
Stem-loop primed reverse transcription does not discriminate between miR-133a-3p and miR-133b-3p. (**A**) Sequence alignments for the miR-1/206 and miR-133 myomiR families. Differences between miRNA family members are highlighted in yellow. Synthetic miRNA mimic oligonucleotide templates were amplified using either the intended on-target small RNA TaqMan assay (i.e. both RT primer and qPCR primer/probe mixes) or the assay for the related off-target miRNA. Amplification plots are shown for (**B**) miR-1a-3p and miR-206-3p, and (**C**) miR-133a-3p and miR-133b-3p. (The threshold is indicated by a dotted line). Diagrams of stem-loop reverse transcription primers in complex with either (**D**) their intended cognate target miRNAs, or (**E**) the unintended off-target related miRNA. High complementarity in the miRNA binding region for the miR-133 family stem-loop primers explains the off-target priming observed for miR-133a-3p and miR-133b-3p.

These data demonstrate that small RNA TaqMan assays are unable to distinguish between miR-133a-3p and miR-133b-3p, and are unlikely to be able to discriminate between other small RNAs which differ only at the 3ʹ terminus in the general case. For this reason, we have chosen not to investigate both miR-133a-3p and miR-133b-3p using RT-qPCR in our studies. Instead, we report only miR-133a-3p data (while acknowledging here that this assay will also amplify the less abundant miR-133b-3p species). Notably, we have previously shown that both miR-133a-3p and miR-133b-3p are upregulated in dystrophic serum using small RNA-seq (which can distinguish between these miRNAs) ^17^.

### Dose-dependent ex-myomiR restoration in *mdx* serum following PPMO treatment

We next sought to analyse the serum levels of putative DMD biomarker myomiRs (miR-1a-3p, miR-133a-3p, miR-206-3p, and miR-483-3p) in the PPMO-treated animals, compared with WT and *mdx* controls. Levels of serum CK were measured in parallel. RNA was extracted from serum samples and miRNA levels determined by small RNA TaqMan RT-qPCR. All miRNA biomarkers were elevated in *mdx* serum (**Figure 3**). The myomiRs were significantly up-regulated in *mdx* serum with median fold-change increases of 16-, 19.9-, and 48-fold for miR-1a-3p, miR-133a-3p, and miR-206-3p, respectively (**Figure 3A-C**). miR-483-3p was up-regulated by 12.5-fold but did not reach statistical significance at the *P*<0.05 level (**Figure 3D**). PPMO-treated *mdx* mice exhibited a dose-dependent shift in all miRNA biomarkers towards wild-type levels. Specifically, the animals in the 12 mg/kg group exhibited biomarker restoration that was close to (and not significantly different from) WT levels, especially in the cases of miR-1a-3p and miR-133a-3p. Conversely, the animals in the 6 mg/kg group exhibited biomarker levels that were intermediate between those of *mdx* and WT animals (**Figure 3A-D**). Highly similar results were observed for serum CK (**Figure 3E**). Notably, the variation in CK measurements was less than that observed for the miRNA biomarkers, and consequently more inter-group statistical significant changes were detected.

**Figure 3.**
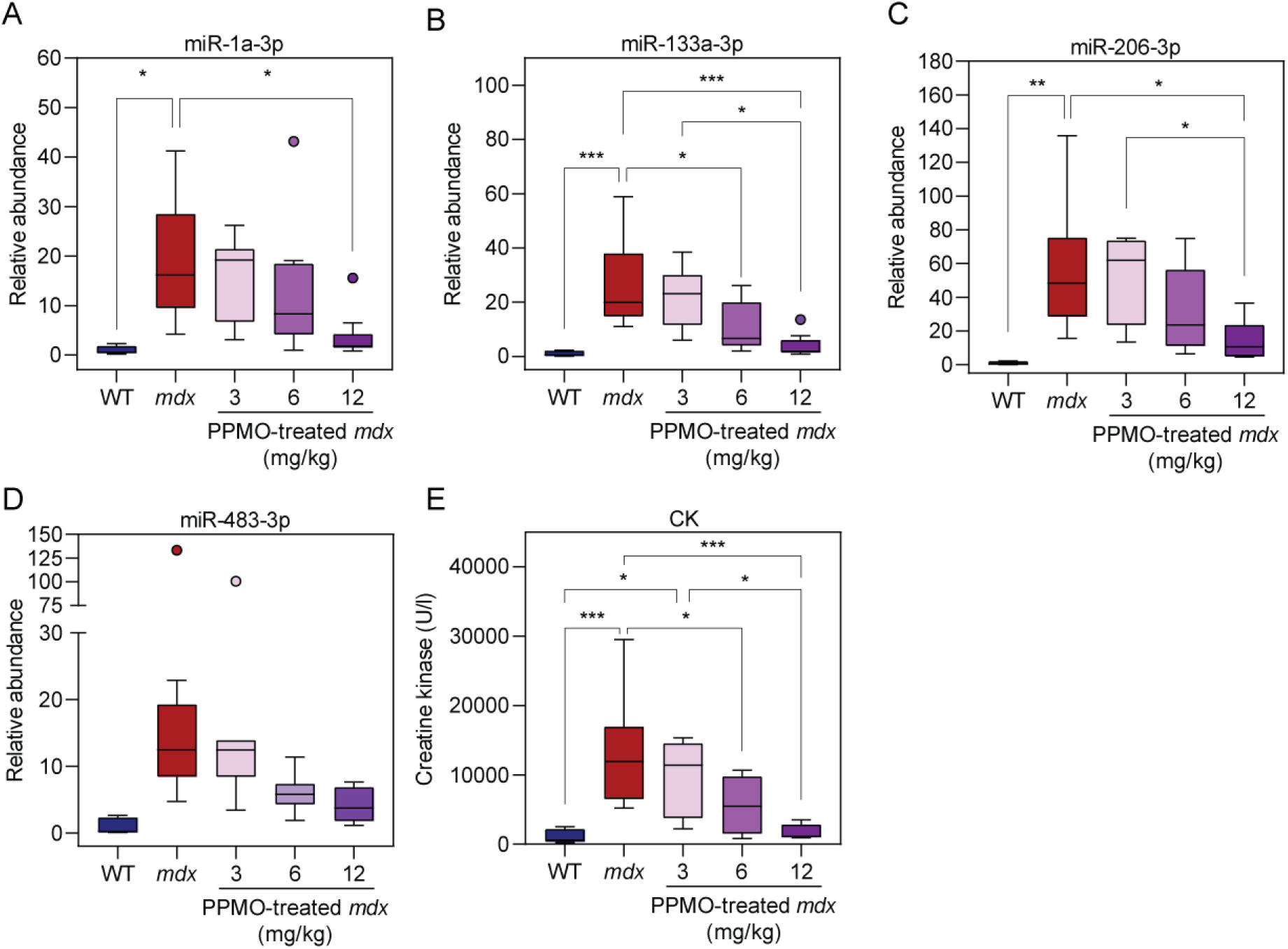
Extracellular myomiRs are pharmacodynamic biomarkers of exon skipping-mediated dystrophin restoration. Eight-week-old male *mdx* mice were treated with a single intravenous injection of PPMO conjugates (Pip9b2-PMO) at one of three doses (3, 6, or 12 mg/kg). Mice were harvested 2 weeks after treatment (age 10 weeks) and serum analysed for (**A**) miR-1a-3p, (**B**) miR-133a-3p, (**C**) miR-206-3p, (**D**) miR-483-3p, and (**E**) creatine kinase (CK). Serum from untreated WT and *mdx* animals served as controls. Data are shown as Tukey box plots, with statistically significant differences tested by one-way ANOVA with Bonferroni *post hoc* test, \**P*<0.05, \*\**P*<0.01, \*\*\**P*<0.001.

### Ex-myomiR biomarkers are anti-correlated with dystrophin expression

Correlation analyses were performed to compare either muscle dystrophin expression or exon skipping levels with matched biomarker levels for PPMO-treated animals (**Figure 4**). Serum biomarker levels were strongly negatively correlated with dystrophin protein expression: miR-1a-3p (*r*=-0.72), miR-133a-3p (*r*=-0.76), miR-206-3p (*r*=-0.64), miR-483-3p (*r*=-0.55), and CK (*r*=-0.75) (all *P*<0.005) (**Figure 4A-E**). Similarly, serum biomarker levels were also strongly negatively correlated with exon 23 skipping: miR-1a-3p (*r*=-0.67), miR-133a-3p (*r*=-0.75), miR-206-3p (*r*=-0.67), miR-483-3p (*r*=-0.64), and CK (*r*=-0.75) (all *P*<0.0006) (**Figure 4F-J**). Conversely, the serum biomarkers were positively correlated with one another (**Figure 4K**, *r*>0.72, *P*<5.2×10^-5^). miR-133a-3p was the most strongly correlated biomarker when compared to dystrophin protein (**Figure 4B**), with serum CK a close second (**Figure 4E**). In contrast, miR-483-3p was the most weakly correlated biomarker (**Figure 4D,I**).

**Figure 4.**
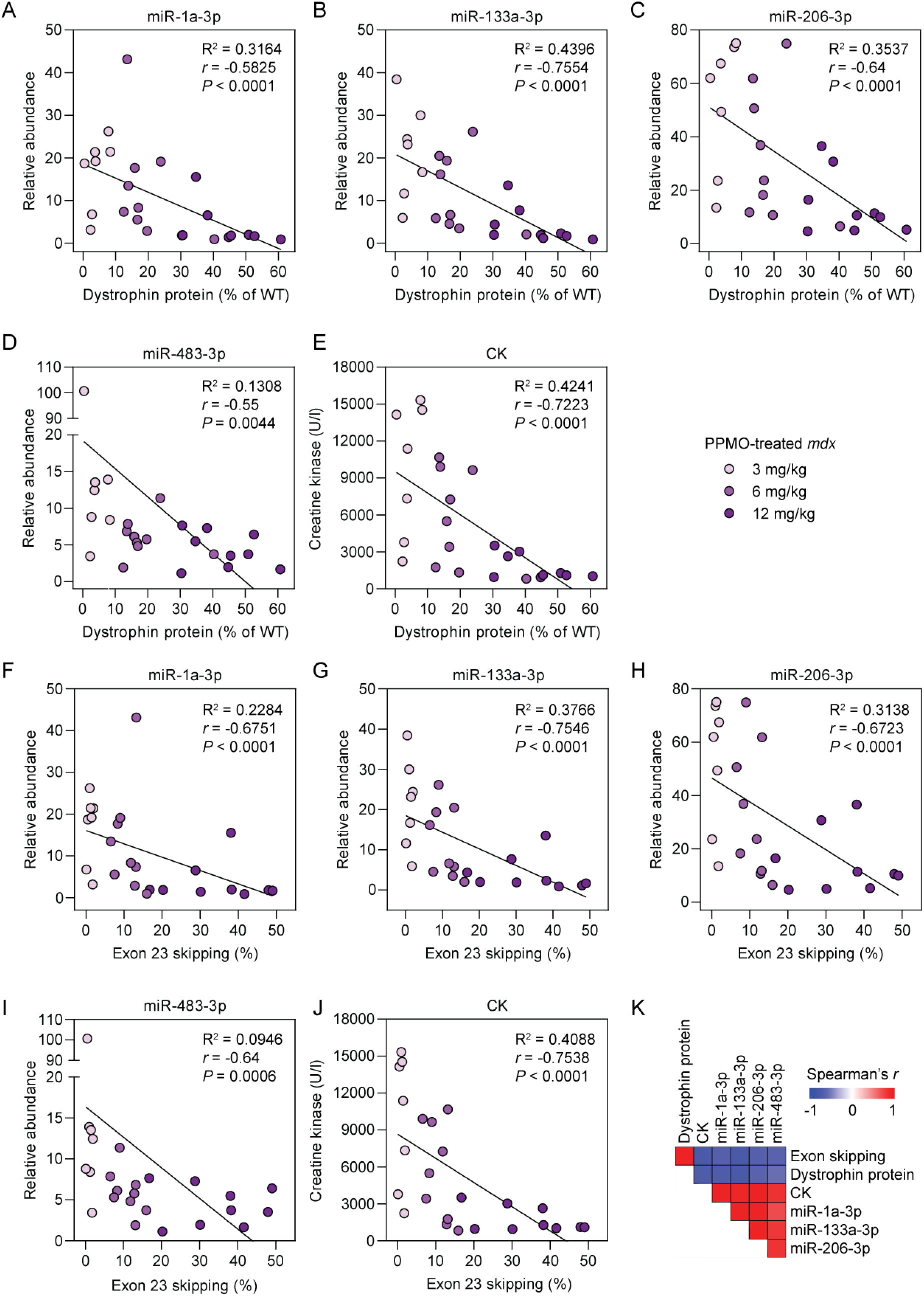
Serum biomarkers are strongly anti-correlated with dystrophin expression and exon 23 skipping levels in PPMO-treated *mdx* mice. Correlation analyses for dystrophin protein expression vs (**A**) miR-1a-3p, (**B**) miR-133a-3p, (**C**) miR-206-3p, (**D**), miR-483-3p, and (**E**) serum CK. Correlation analyses for *Dmd* exon 23 skipping vs (**F**) miR-1a-3p, (**G**) miR-133a-3p, (**H**) miR-206-3p, (**I**), miR-483-3p, and (**J**) serum CK. Values were analysed using linear regression and Spearman correlation analyses. (**K**) Heatmap of Spearman correlation coefficients for all pair-wise comparisons. Red indicates a positive correlation and blue indicates a negative correlation.

### Dose-independent uniform dystrophin distribution after PPMO treatment

We previously showed that dystrophin distribution is uniform after treatment with a single 12.5 mg/kg dose of Pip9b2-PMO in the *mdx* mouse ^9^. This is in contrast to the patchy pattern of dystrophin expression observed in *mdx-Xist*^Δhs^ mice expressing equivalent total levels of dystrophin as that achieved with exon skipping ^9^. We were therefore motivated to assess whether a within-fiber uniform pattern of dystrophin expression is observed at all PPMO doses tested. To this end, we analysed dystrophin expression by immunofluorescence staining in TA muscle sections in both transverse and longitudinal orientations (**Figure 5**). PPMO-treated *mdx* mice generally exhibited a uniform pattern of dystrophin expression. In the low dose (3 mg/kg) group, some degree of between-fiber patchiness was observed when sections were viewed in transverse orientation (**Figure 5**). Specifically, some fibers were dystrophin-positive, whereas others were dystrophin negative. The positively-stained fibers generally exhibited consistent staining around the circumference of the fiber, although the strength of the staining was also weak in some fibers. In contrast, within-fiber patchiness was not observed, as viewed in longitudinal orientation (**Figure 5**).

**Figure 5.**
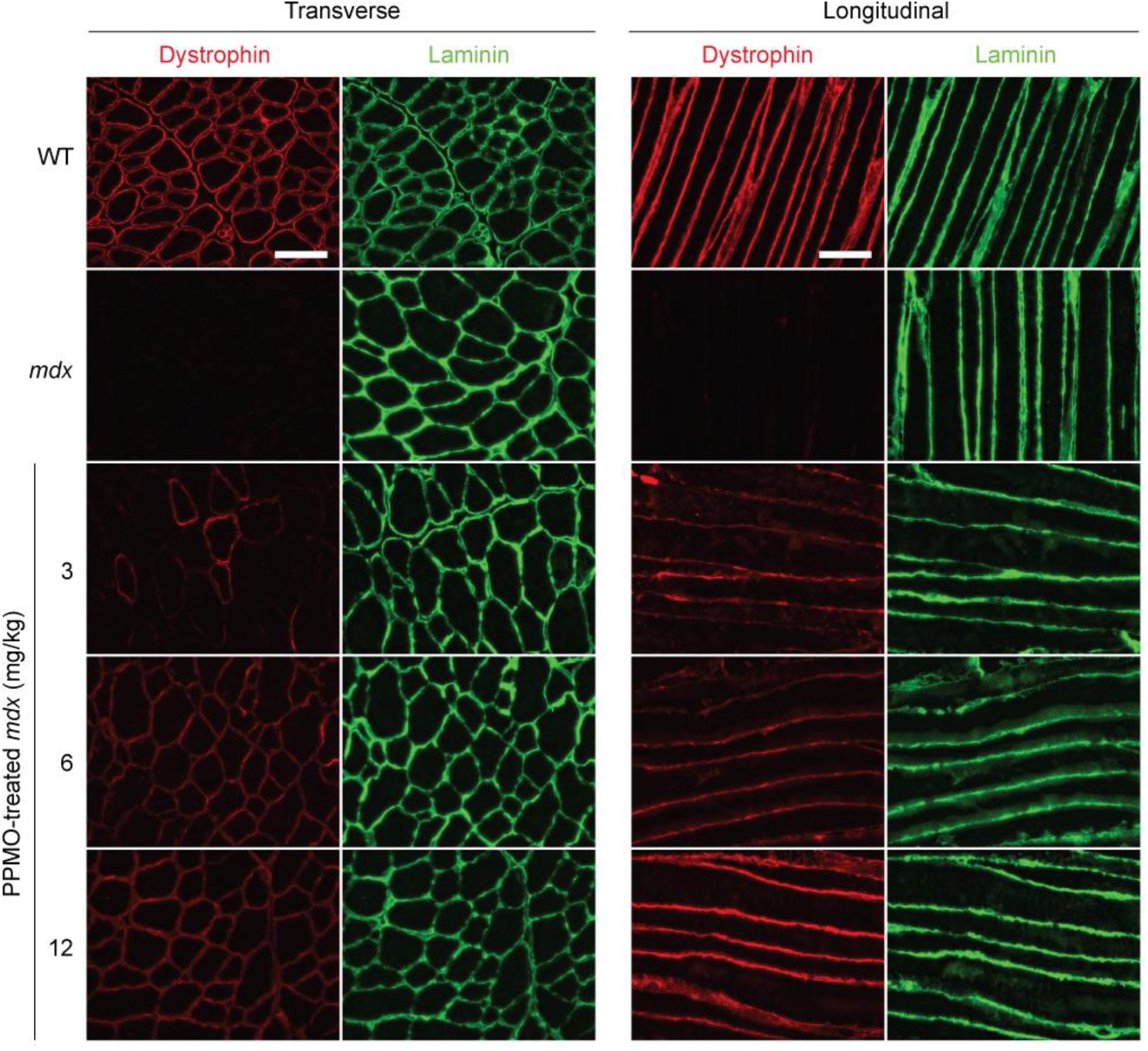
Distribution of sarcolemmal dystrophin expression in PPMO-treated *mdx* mice. Representative dystrophin immunofluorescence images of TA muscle sections in both transverse and longitudinal orientations for *mdx* mice treated with a single dose of PPMO (at 3, 6, or 12 mg/kg doses). Untreated WT and *mdx* samples were analysed in parallel as controls. Sections were co-strained for laminin to delineate myofiber boundaries. Images were taken at 20× magnification, and scale bars represent 100 μm.

### Dose-dependent restoration of myofiber elasticity following PPMO treatment

A subset of the mice described above was further analysed using by atomic force microscopy (AFM) nanoindentation assay in order to investigate the relationship between dystrophin restoration, circulating biomarker levels, and the biophysical properties of muscle. To this end, TA muscle explants (all *n*=4) from WT, *mdx*, and PPMO-treated *mdx* mice at the three doses: 3 mg/kg, 6 mg/kg, or 12 mg/kg, and were assessed by AFM. Force versus displacement measurements were performed and Young’s modulus (*E*) (a measure of the stiffness of the biological sample in response to an applied load) calculated as described previously ^42^. An indentation depth of 1050 nm was utilized such that Young’s modulus measurements reflect the stiffness of the myofiber inner structure, deeper than the actin microfilaments directly underlying the sarcolemma.

The mean *E* values were 4.7 kPa and 1.7 kPa for the WT and *mdx* mice, respectively (a 2.7-fold reduction in *mdx* muscle, *P*<0.0001), indicative of markedly lower resistance of dystrophic muscle to reversible deformation (**Figure 6A**), consistent with previous observations ^42,43^. Treatment with PPMO resulted in a dose-dependent shift in the mean *E* values (**Figure 6A**). Young’s modulus values were not significantly different between *mdx* mice treated with PPMO at the lowest dose, i.e. 3 mg/kg and untreated *mdx* mice (**Figure 6A**), whereas the 6 mg/kg PPMO treatment was sufficient to completely reverse the reduction in stiffness observed in *mdx* muscle (**Figure 6A**). Interestingly, treatment with 12 mg/kg PPMO resulted in a ∼2-fold increase (*P*<0.0001) in Young’s modulus (mean *E*=9.2 kPa) relative to the WT control group, suggesting a higher resistance to the deformation as compared to that in the control mice (**Figure 6A**). Very similar results were observed using measurement depths of 250 and 550 nm (**Figure S3**).

**Figure 6.**
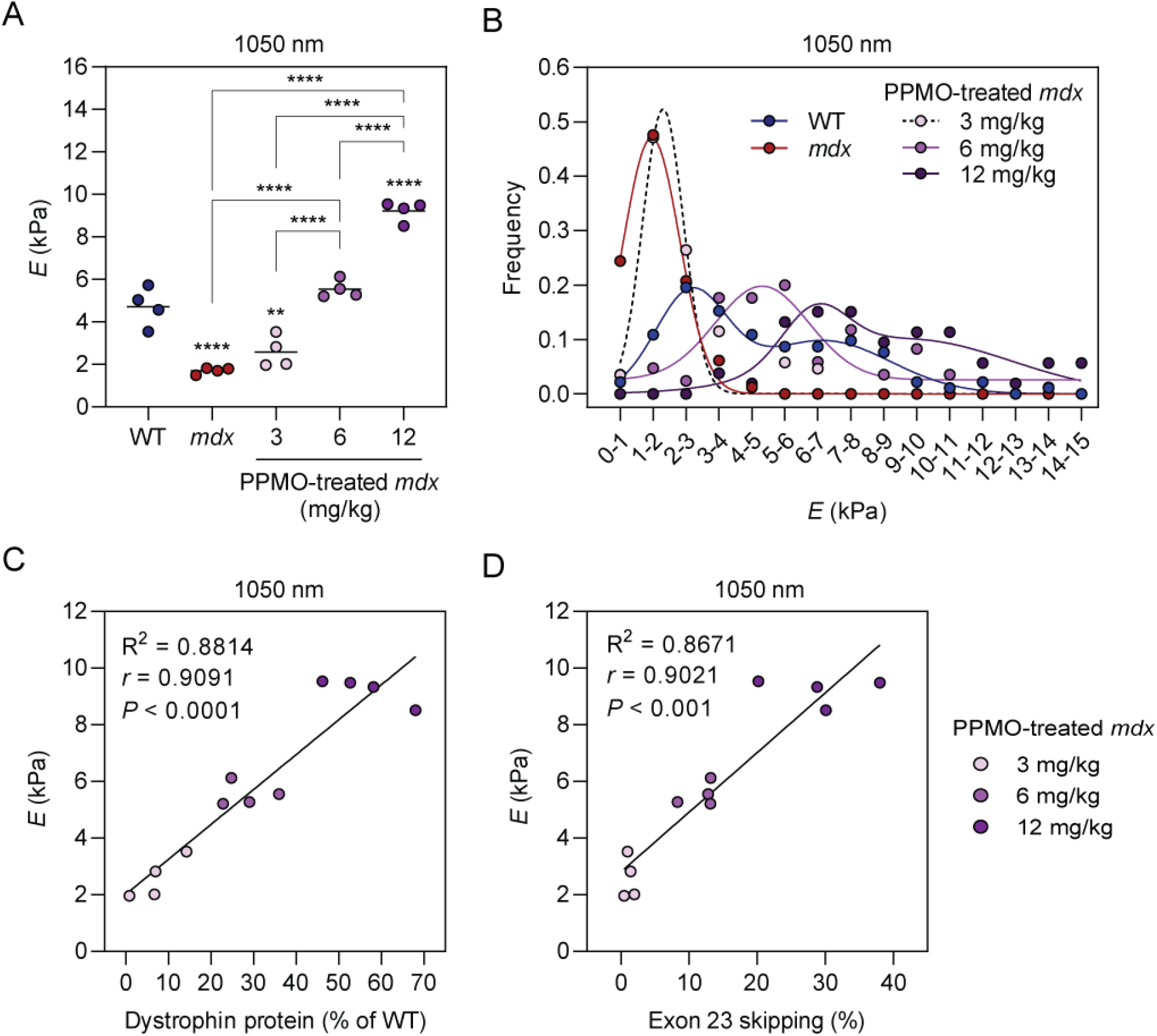
Dose-dependent changes in muscle stiffness following exon skipping-mediated dystrophin rescue. Atomic force microscopy was performed in order to determine Young’s modulus (*E*) calculated for an indentation depth of 1050 nm in TA muscle explants from WT, *mdx*, and PPMO-treated *mdx* mice and plotted as (**A**) mean *E* values for each individual sample (representing 60 to 100 elasticity maps, depending on the sample group), and (**B**) the distribution of all determined *E* values. Mean values and individual sample data points are shown. Statistical significance was tested by one-way ANOVA and Bonferroni *post hoc* test. Statistical comparisons are made to the WT control group unless otherwise indicated, \*\**P*<0.01, \*\*\*\**P*<0.0001. Correlation analyses for Young’s modulus vs (**C**) dystrophin protein expression, and (**D**) *Dmd* exon 23 skipping in PPMO-treated *mdx* samples. Values were analysed using linear regression and Spearman correlation analyses.

Inspection of underlying *E* value determinations revealed a Gaussian distribution in *mdx* muscle with an *E* maximum at 1-2 kPa, consistent with high myofiber elasticity (**Figure 6B**). WT *E* values exhibited a bimodal distribution with maxima at 2-3 kPa and 7-8 kPa, (**Figure 6B**). The PPMO-treated *mdx* samples showed a gradation of effect, whereby the distribution of *E* values shifted from an *mdx*-like distribution in the 3 mg/kg group, to a more WT-like distribution in the 12 mg/kg group. For the low dose 3 mg/kg group, the data were fit to a Gaussian distribution similar to the *mdx* samples, although a smaller population of measurements with *E* values ∼5-7 kPa was also observed, which did not fit the calculated distribution. The results obtained for 6 mg/kg and 12 mg/kg data were best fitted with the sum of two Gaussian distributions, although the 6 mg/kg data were monomodal, and the 12 mg/kg data were bimodal. The distribution of *E* values in the high dose group resembled that of WT muscle, although the *E* values were shifted to the right (with a maximum at 6-8 kPa) and a high proportion of stiffer measurements (including up to 14-15 kPa), indicating an increase in myofiber stiffness in the PPMO-treated samples relative to WT (**Figure 6B**). Mean Young’s modulus was strongly positively correlated with both dystrophin protein expression (*r*=0.909, *P*<0.0001) (**Figure 6C**), and exon 23 skipping (*r*=0.902, *P*<0.01) (**Figure 6D**) in PPMO-treated *mdx* mice. Overall, these findings suggest that exon skipping-mediated dystrophin rescue can restore the biophysical properties of dystrophic muscle.

## Discussion

Here we show that treatment of dystrophin-deficient *mdx* mice with Pip9b2-PMO results in dose-dependent *Dmd* exon 23 skipping, dystrophin rescue, restoration of circulating miRNA biomarker levels, and correction of muscle biophysical properties. Treatment of *mdx* mice with PPMO conjugates resulted in a within-fiber uniform pattern of sarcolemmal dystrophin expression across dose levels (**Figure 5**), consistent with our previous results ^9^. This is in contrast with our observations in CRISPR-Cas9-treated muscle (**see related manuscript**), whereby dystrophin was expressed in a patchy manner. These findings point to a potential advantage of antisense oligonucleotide-mediated exon skipping over gene editing ^44^.

This study adds to the growing body of literature that supports the use of extracellular myomiRs as minimally-invasive biomarkers for use in DMD patients. Notably, the negative correlations between circulating miRNA levels and molecular correction of dystrophin (**Figure 4**) suggests that these miRNAs could be utilized as pharmacodynamic biomarkers to assess the effectiveness of experimental therapies in clinical trials, or approved therapies in patients.

These findings are consistent with observations from a previous study from our group whereby two PPMO conjugates with differing potencies where found to induce corresponding levels of biomarker restoration ^20^. Importantly, several lines of evidence indicate that ex-myomiRs are not biomarkers of dystrophin expression *per se*. Firstly, the elevation of serum myomiR levels (i.e. miR-1, miR-133, and miR-206) has been reported in the context of multiple other myopathies ^27,37,45,46^, and so is not a unique feature of dystrophinopathy. Secondly, serum myomiR biomarkers have been shown to be restored to wild-type levels in DMD mouse models following experimental therapies that do not involve dystrophin restoration. Specifically, serum miR-206 was restored to wild-type levels in *mdx* mice treated with a myostatin pathway blockade strategy (i.e. in the context of no dystrophin restoration) ^47^, and serum myomiRs are also present at wild-type levels in *mdx*-Fiona mice (which lack dystrophin but overexpress the dystrophin-paralogue, utrophin) ^48^. Instead, we have suggested that ex-myomiR levels are reflective of muscle turnover (i.e. myonecrosis and compensatory regeneration) based on several lines of evidence: (i) ex-myomiRs are elevated during perinatal myogenesis in both WT and *mdx* animals ^32^, (ii) myomiRs are progressively released by human and murine myoblasts cultures undergoing differentiation ^32^, and (iii) a biphasic pattern of ex-myomiR release was observed in *mdx* mice following an acute exercise regimen, corresponding to an initial damage phase immediately after exercise, and a secondary regenerative phase 5-7 days post exercise^32^.

Analysis of the biophysical properties of muscle using an AFM nanoindentation assay revealed reduced resistance to mechanical deformation (i.e. stiffness) in dystrophic muscle (**Figure 6**), consistent with previous reports ^42,43^. PPMO treatment restored myofiber elastic properties in a dose-responsive manner, although the high PPMO dose increased stiffness beyond that observed in WT mice. The reason for this is unclear, but given that treated *mdx* mice exhibit signs of muscle hypertrophy ^9^ (**Figure 5**), the resulting increase in the concentration of protein components of the contractile apparatus and cytoskeletal proteins may contribute to this increase in myofiber stiffness. Additionally, the myonuclei of *mdx* myofibers are often arranged in centrally-nucleated chains, indicative of recent regeneration events, which may also contribute to an increase in stiffness ^49^. These data provide further evidence that serum miRNA measurements can be used to provide information about the underlying muscle (patho)physiology. Similarly, AFM has been used to assess the elastic properties of DMD patient biopsy material, suggesting that this technique could be utilized for diagnostic purposes ^42^. The data contained in this study provide evidence that AFM might also be suitable for assessing the effectiveness of therapeutic interventions for DMD (**Figure 6**), although invasive biopsy collection is still required.

Notably, in this study serum CK arguably performed better than the ex-myomiRs. CK is known to be highly variable in patients and affected by a number of confounding factors (e.g. exercise ^50,51^, age ^52^, race ^53^, and drugs such as statins ^54^), which may not impact serum myomiR levels, as has been reported by others ^18^. Furthermore, we previously observed that serum myomiR levels in human serum provided very high predictive power for distinguishing between DMD and healthy individuals (AUC>0.99 for miR-206-3p, *n*=28/16 DMD patients/healthy controls) ^17^. An important consideration is that the measurement of serum miRNA biomarkers requires multiple processing steps (i.e. RNA extraction, reverse transcription, qPCR amplification) whereas serum CK can be measured directly by either ELISA or enzymatic activity assay. As such, serum miRNA measurements may be subject to a greater degree of technical variation, especially in a research laboratory setting.

In this study, we also aimed to address the suitability of the small RNA TaqMan RT-qPCR method to distinguish between closely-related myomiRs. This commercial technology has been extensively used in the study of miRNAs across many areas of biology and medicine, and was shown to provide excellent discrimination between members of the let-7 family of miRNAs (some of which differ by only a single nucleotide, which is not the 3ʹ terminus) ^41^. However, our data clearly show that this product is incapable of distinguishing between miRNAs that differ only in their 3ʹ terminal nucleotide, such as miR-133a-3p and miR-133b-3p, and that the lack of discrimination occurs at the RT step (**Figure 2**). We note that a similar failure to distinguish between miR-133a and miR-133b was reported by Ikeda *et al*. although the exact method, and procedures used are not clear ^55^. This technical point is important, given that multiple studies have reported expression data for both of these miRNAs using this technology ^23,29,34,35,37,38,40^. The present study underlines the importance of assay validation, and how it should not be assumed that commercial products are valid *a priori*.

In conclusion, we show that PPMO-mediated exon skipping results in dose-dependent dystrophin rescue and restoration of the biophysical properties of dystrophic muscle. Importantly, a within-fiber uniform pattern of dystrophin expression was observed at all dose levels. Serum myomiR levels were inversely correlated with dystrophin expression and myofiber stiffness, thereby providing support for their use as pharmacodynamic biomarkers in the context of DMD.

## Materials and Methods

### Animal Studies

All procedures were authorized by the UK Home Office (project license 30/2907) in accordance with the Animals (Scientific Procedures) Act 1986. Animals were housed with a 12:12 hour light:dark cycle, with access to food and water *ad libitum*. Mice used were male wild-type C57 (C57BL/10ScSn) or dystrophic *mdx* (C57BL/10ScSn-*Dmd^mdx^*/J).

### PPMO

PPMO conjugates consisted of a peptide (Pip9b2, Ac-RXRRBRRFQILYRBRXRB-OH, where ‘X’ is aminohexanoic acid, and ‘B’ is β-alanine) covalently attached to a PMO designed to induce skipping of *Dmd* exon 23 (**Table S1**). The peptide was synthesized by standard Fmoc solid phase chemistry, and conjugated to the PMO via the amine of the morpholine heterocycle at its 3ʹ terminus, as described previously ^56^. Conjugates were dissolved in sterile water and passed through 0.22 µm cellulose acetate filters.

Male *mdx* mice were injected with a single dose of PPMO conjugates (either 3, 6, or 12 mg/kg) in 0.9% sterile saline via the tail vein at 8 weeks of age. Animals were sacrificed two weeks later (at 10 weeks of age) by escalating CO_2_ concentration, and serum/muscle tissues harvested.

TA muscles were macrodissected and flash-frozen in isopentane cooled on dry ice. Muscles were mounted on corks using Tissue-TEK O.C.T. Compound (VWR, Lutterworth, UK) to enable cryosectioning, and stored at -80°C.

Blood was harvested from the jugular vein and collected using Microvette CB 300 collection tubes (Sarstedt, Nümbrecht, Germany) as described previously ^57^. Blood was incubated at 4°C for at least 2 hours to facilitate clotting and then cellular blood pelleted by centrifugation at 10,000 *g* for 5 minutes. The serum supernatant was transferred to a fresh microcentrifuge tube and samples stored at -80°C until ready for analysis.

### Western Blot

TA muscles were homogenized in modified RIPA buffer (50 mM Tris pH8, 150 mM NaCl, 1% IGEPAL CA-630, 0.5% sodium deoxycholate, 10% SDS and 1× cOmplete protease inhibitor cocktail (Merck, Feltham, UK)) using a Precellys 24 Tissue Homogenizer (Bertin Instruments, Montigny-le-Bretonneux, France) (4 cycles at 5,000 rpm for 30 seconds). 20 µg of protein was mixed with NuPAGE sample reducing agent and NuPAGE LDS sample buffer and run on NuPAGE 3 to 8% Tris-Acetate gel (all Thermo Fisher Scientific, Abingdon, UK). Protein was electrotransferred onto a 0.45 µm polyvinylidene difluoride (PVDF) membrane for 1 hour at 30 V, followed by 1 hour at 100 V and blocked with Odyssey blocking buffer (LI-COR Biosciences Ltd, Cambridge, UK) for 1 hour at room temperature. Membranes were incubated with primary antibodies (**Table S2**) overnight in Odyssey blocking buffer supplemented with 0.1% Tween-20 at 4°C. Membranes were washed 3 times with PBST and then incubated with HRP-conjugated secondary antibodies (**Table S2**) in Odyssey blocking buffer supplemented with 0.1% Tween-20 at room temperature for 1 hour. Chemiluminescent signal was detected using Clarity Western ECL substrate (Bio-Rad, Watford, UK).

### Exon 23 skipping RT-qPCR

Tissues were homogenized using a Precellys homogenizer (Bertin Instruments) (2 cycles at 5,500 rpm for 30 seconds), and RNA was extracted using TRIzol Reagent (Thermo Fisher Scientific) according to manufacturer’s instructions. cDNA was generated from 500 ng of total RNA using the High-Capacity cDNA Reverse Transcription Kit (Thermo Fisher Scientific) according to manufacturer’s instructions. The resulting cDNA was diluted 1:5 prior to analysis. cDNA was amplified using a StepOne Plus real-time PCR Thermocycler with TaqMan Gene Expression Master Mix (both Thermo Fisher Scientific) using universal cycling conditions: 95°C for 10 min, followed by 45 cycles of 95°C for 15 seconds, 60°C for 1 minute. All samples were analysed in duplicate. Absolute quantification was performed by comparing samples to standard curves comprised of serial dilutions of the full length (unskipped) and Δ23 *Dmd* (skipped) DNA target templates (IDT, Leuven, Belgium). The degree of exon 23 skipping was determined by calculating the percentage of skipped transcripts relative to the total (i.e. skipped + unskipped). Sequences of RT-qPCR primers and probes are shown in **Table S3**.

### Immunofluorescence

Fresh frozen TA muscles were mounted onto corks with Tissue-TEK O.C.T. Compound and cryosectioned (8 µm) in transverse and longitudinal orientations. Sections were transferred onto SuperFrost Plus microslides (VWR), left to dry for 10 minutes at room temperature and stored at -80°C until ready for analysis. Slides were air-dried, soaked in PBS for 10 minutes at room temperature, and then blocked in PBS supplemented with 20% foetal calf serum (FCS, Thermo Fisher Scientific) and 20% normal goat serum (NGS, MP Biomedicals, Eschwege, Germany) for 2 hours at room temperature. Subsequently, slides were incubated with primary antibodies in PBS supplemented with 20% FCS and 20% NGS for 2 hours at room temperature. After washing three times with PBS, slides were incubated with fluorophore-conjugated secondary antibodies in PBS for 1 hour at room temperature. Slides were then washed three times with PBS, and mounted using Dako, Fluorescence Mounting Medium (Agilent, Didcot, UK). Details of all antibodies are shown in **Table S2**.

### Serum Biomarker Analysis

Serum miRNA analysis was performed as described briefly ^57,58^. Briefly, RNA was extracted from 50 µl of blood serum using TRIzol LS (Thermo Fisher Scientific) according to manufacturer’s instructions with minor modifications. A synthetic spike-in control oligonucleotide (i.e. cel-miR-39, 2.5 fmol, **Table S1**) was added at the phenolic extraction phase, and mixed thoroughly. Additionally, 150 µl nuclease-free water (Thermo Fisher Scientific) was added at the phenolic extraction phase in order to increase the volume of the aqueous phase after phase separation. RNA was precipitated using isopropanol, and 1 µl of RNase-free glycogen (Roche, Welwyn Garden City, UK) was added to each sample as an inert carrier to assist RNA recovery. Pellets were air-dried, and then RNA resuspended in 30 µl nuclease-free water, and samples incubated at 55°C for 10 minutes. Samples were stored at -80°C until ready for analysis. Reverse transcription was performed with the TaqMan MicroRNA Reverse Transcription Kit (Thermo Fisher Scientific) as according to manufacturer’s instructions using 10 µl of each RNA sample. Reverse transcription reactions were performed with miRNA-specific stem loop primers as appropriate. The resulting reverse transcription product was diluted 1:2 prior to analysis, and 2 µl of cDNA was added per 20 µl qPCR reaction. cDNA was amplified as described above. Data were normalized to cel-miR-39 ^58^ and data analyzed using the Pfaffl method ^59^. Details of small RNA TaqMan assays are provided in **Table S4**.

To validate the specificity of miRNA RT-qPCR assays, 50 fmol solutions containing synthetic oligonucleotide mimics of each miRNA (**Table S1**, IDT) were analysed using the correct RT primer or related off-target RT primer, and the resulting cDNA analysed using the various qPCR assays.

Serum CK analysis was performed as a service at the MRC Harwell Institute (Oxford, UK).

### Atomic Force Microscopy Nanoindentation Assay

Mouse muscles were stored under liquid nitrogen in DMEM supplemented with 10% DMSO following the protocol published elsewhere ^42^. One hour before AFM measurements, muscles were unfrozen at room temperature by placing them in a Petri dish filled with DMEM (Sigma-Aldrich, Burlington, MA, USA). Afterwards, they were washed twice in DMEM (for 2 minutes each time). The muscles were next removed from the medium, and the bottom part was gently dried with filter paper, followed by gluing them on to a glass coverslip using two 0.5 µl droplets of cyanoadhesive placed at both extremities as described previously ^43^. The muscle sample was immediately immersed in DMEM, and AFM measurements were performed (no longer than 3 hours per individual muscle).

Measurements of the mechanical properties of muscles were carried out using an atomic force microscope (AFM, Xe120, Park Systems, Korea) equipped with a liquid cell sitting on a piezoelectric scanner with an *XY* range of 100 µm × 100 µm. AFM worked in a force spectroscopy mode that allows recording an approach and retracting the AFM probe (i.e. cantilever) using a separate piezoelectric scanner with a Z-range of 25 µm. Cantilevers were silicon nitride cantilevers of the ORC-8 (Bruker) type characterized by a nominal resonant frequency of 18 kHz, open-angle of 36°, and nominal spring constant of 0.1 N/m. The spring constant of the cantilevers was calibrated using the Sader method ^60^. It ranges from 0.101 N/m to 0.110 N/m. Force curves, i.e. dependencies between cantilever deflection and relative sample position, were recorded to estimate the elastic properties of muscles. Force curves were recorded in 36 positions (a grid of 6 pixels × 6 pixels was set). For each muscle sample, 28-31 maps were recorded (in total, 1008 to 1116 force curves were analyzed). Measurements were acquired at an approach and retract speed of 8 µm/s. Calibration curves (curves recorded on a stiff, non-deformable surface) were collected from the measurements of a silicon nitride rectangular chip glued at the muscle height level.

The contact mechanics model analyses the relationship between the load force and indentation. In the AFM, to obtain indentation values, force curves were recorded on a stiff, non-deformable surface (here, glass coverslip) were subtracted from force curves collected on a muscle sample. The resulting force versus indentation curve were fitted with the Hertz-Sneddon contact mechanics assuming that the indenting AFM probe can be modelled as a cone ^61^. For such a probe, the relation between the load force *F* and the resulting indentation depth *δ* is:

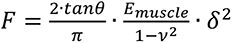

where *Θ* is the open-angle of the probing cone (here, 36°), *ν* is the Poisson’s ratio defining the compressibility of the studied material (here, *ν=*0.5 because cells are incompressible), and *E_muscle_* is the Young’s (elastic) modulus of the studied muscle sample. Young’s modulus was determined at three indentation depths, i.e. 250 nm, 500 nm, and 1050 nm.

### Statistics

Statistical analyses were performed using GraphPad Prism v9 (GraphPad Software Inc, La Jolla, CA). Differences between groups were compared using one-way ANOVA and Bonferroni *post hoc* test. Differences were considered significant at the *P*<0.05 level. Other analyses performed in GraphPad were the Shapiro-Wilk and Kolmogorov-Smirnov tests for normality, linear regression, non-linear regression (Gaussian and sum of two Gaussian), and Spearman correlation analysis was performed. Data were tested for outliers using Grubb’s test, after which one sample in the 12 mg/kg group (which exhibited extremely high biomarkers levels) was removed from all analyses. Heatmaps were produced using TMeV (The Institute for Genomic Research, Rockville, MD, USA) ^62^.

## Acknowledgements

K.C. is supported by a studentship from the Clarendon Fund. Work in the laboratory of MJAW is supported by grants from the UK Medical Research Council.

