## Supplementary Information for "PPMO-mediated exon skipping induces uniform sarcolemmal dystrophin rescue with dose-dependent restoration of circulating microRNA biomarkers and muscle biophysical properties"

| ID | Sequence (5' to 3') |
|---|---|
| **Exon skipping oligonucleotides (PMO)** | |
| ***Dmd* Ex23 skipping** | GGCCAAACCTCGGCTTACCTGAAAT |
| **miRNA mimics/spikes (RNA)** | |
| **mmu-miR-1a-3p** | UGGAAUGUAAAGAAGUAUGUAU |
| **mmu-miR-133a-3p** | UUUGGUCCCCUUCAACCAGCUG |
| **mmu-miR-133b-3p** | UUUGGUCCCCUUCAACCAGCUA |
| **mmu-miR-206-3p** | UGGAAUGUAAGGAAGUGUGUGG |
| **cel-miR-39** | UCACCGGGUGUAAAUCAGCUUG |

**Table S1**

**Oligonucleotides used in this study.**

| Target protein | Host | Product ID | Manufacturer | Dilution |
|---|---|---|---|---|
| **Dystrophin (DMD)** | Mouse mAb | NCL-DYS1 | Leica Biosystems | 1:100 |
| **Vinculin (VCL)** | Mouse mAb | V9131 | Sigma-Aldrich/Merck | 1:200 |
| **Anti-mouse IgG, HRP-linked** | Horse | 7076 | Cell Signaling Technology | 1:10,000 |
| **Dystrophin (DMD)** | Rabbit pAb | ab15277 | Abcam | 1:1,000 |
| **Laminin subunit alpha 2 (LAMA2)** | Rat mAb | L0663 (clone: 4H8-2) | Sigma-Aldrich/Merck | 1:1,000 |
| **Anti-rabbit IgG Alexa Fluor 594** | Goat | ab150080 | Abcam | 1:500 |
| **Anti-rat IgG Alexa Fluor 488** | Goat | ab150157 | Abcam | 1:500 |

**Table S2**

**Antibodies used in this study.**

| ID | Sequence (5' to 3') |
|---|---|
| **Exon Skipping RT-qPCR** | |
| **qExon22-24-Fwd** | CTGAATATGAAATAATGGAGGAGAGACTCG |
| **qExon22-24-Rev** | CTTCAGCCATCCATTTCTGTAAGGT |
| **qExon22-24-Probe** | /5FAM/ATGTGATTC/ZEN/TGTAATTTCC/3IABkFQ/ |
| **qExon23-24-Fwd** | CAGGCCATTCCTCTTTCAGG |
| **qExon23-24-Rev** | GAAACTTTCCTCCCAGTTGGT |
| **qExon23-24-Probe** | /5HEX/TCAACTTCA/ZEN/GCCATCCATTTCTGTAAGGT/3IABkFQ/ |

**Table S3**

**List of primer sequences used in this study.**

Exon skipping RT-qPCR assay probes include 5′ terminal fluorophores (either FAM or HEX), 3′-terminal Iowa Black fluorescence quencher, and contain internal ZEN modifications. Primer probe assays were obtained from IDT.

| Target | Product ID | Detection Channel |
|---|---|---|
| **Small RNA TaqMan RT-qPCR** | | |
| mmu-miR-1a-3p | 002222 | FAM |
| mmu-miR-133a-3p | 002246 | FAM |
| mmu-miR-206-3p | 000510 | FAM |
| mmu-miR-483-3p | 002560 | FAM |
| cel-miR-39 | 000200 | FAM |

**Table S4**

**List of TaqMan assays used in this study.**

All products were obtained from Thermo Fisher Scientific.

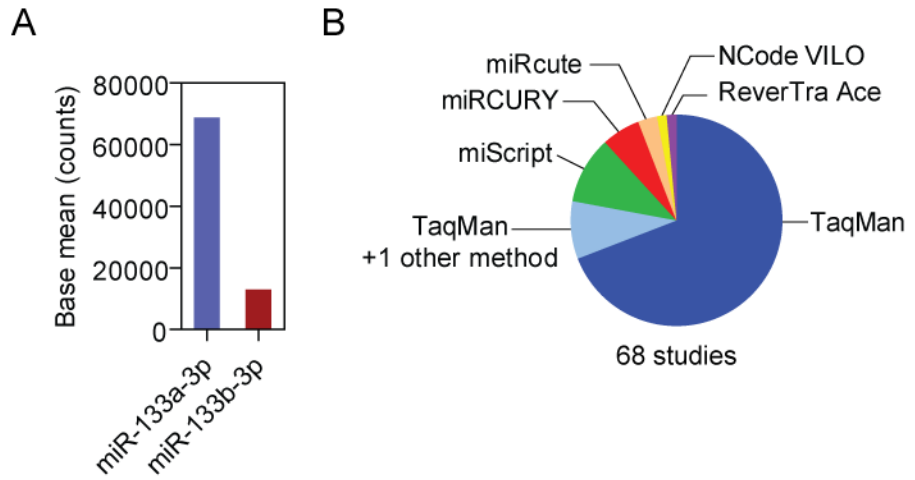


**Figure S1**

**Analysis of miR-133a/b expression and miRNA detection method usage.**

(**A**) Small RNA-seq data for mouse muscle (WT, *mdx*, and PPMO-treated *mdx*). Base mean values (i.e. the average across all samples) are shown. These data show that miR-133b-3p is expressed at much lower levels than miR-133a-3p. Full dataset was published as Coenen-Stass *et al.* [17] and are publicly available (SRA project identifier: SRP102619). (**B**) Analysis of miRNA RT-qPCR methodologies utilized in 68 studies related to miRNA biomarkers in DMD, other myopathies, and muscle biology more generally. The Small RNA TaqMan method is by far the most commonly used technology. Full details of this analysis are in **File S1**.

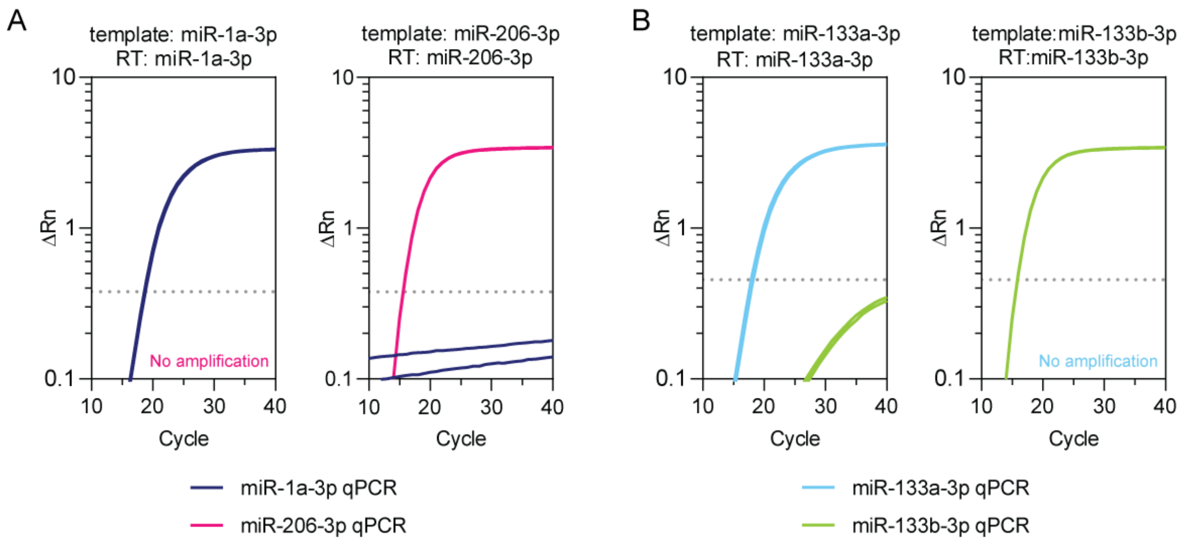


**Figure S2**

**Small RNA TaqMan assays can discriminate between closely-related cDNA templates.**

Synthetic miRNA mimic oligonucleotide templates were reverse transcribed using the correct RT primer, and then amplified using either the on-target TaqMan assay or the off-target TaqMan assay for the closely-related miRNA. Amplification plots are shown for (**A**) miR-1a-3p and miR-206-3p, and (**B**) miR-133a-3p and miR-133b-3p. The threshold is indicated by a dotted line. When reactions were performed in this manner, there was either no amplification for the off-target assays or the amplification curves failed to cross the threshold within 40 cycles.

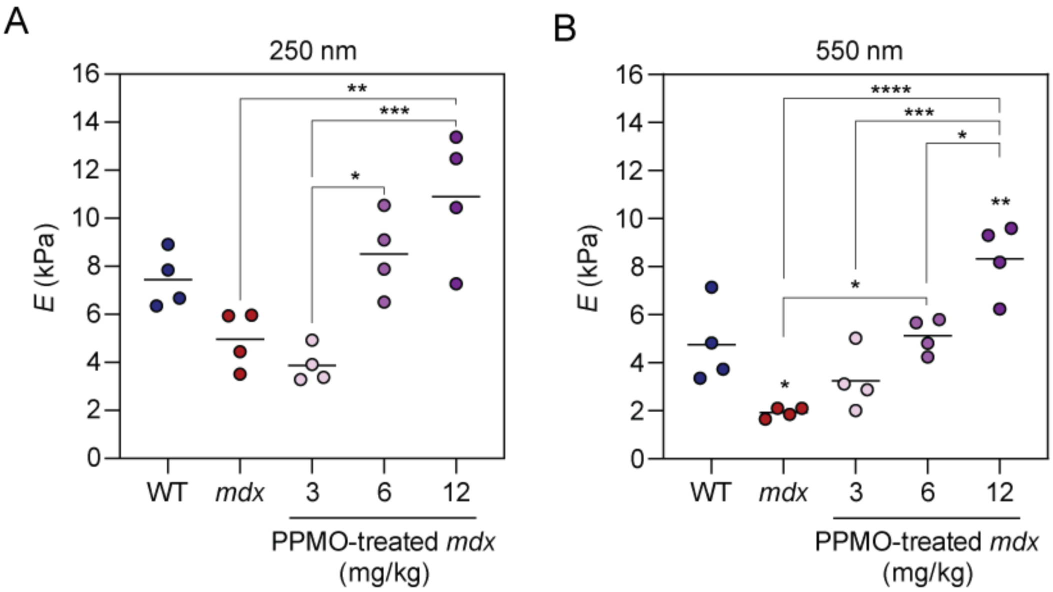


**Figure S3**

**Dose-dependent changes in muscle stiffness following exon skipping treatment.**

Atomic force microscopy was performed in order to determine Young's modulus (*E*) calculated for indentation depth depths of (**A**) 250 nm and (**B**) 550 nm, in TA muscle explants from WT, *mdx*, and PPMO-treated *mdx* mice. Mean values and individual sample data points are shown. Statistical significance was tested by one-way ANOVA and Bonferroni *post hoc* test. Statistical comparisons are made to the WT control group unless otherwise indicated, *$P<0.05$, **$P<0.01$, ***$P<0.001$, ****$P<0.0001$.
